# Lipid packing and local geometry influence septin curvature sensing

**DOI:** 10.1101/2025.02.12.637894

**Authors:** Brandy N. Curtis, Ellysa J. D. Vogt, Christopher Edelmaier, Amy S. Gladfelter

## Abstract

Septins assemble into scaffolds that direct cell growth and morphology that are often localized to the plasma membrane. While septins preferentially bind convex membranes via amphipathic helices, their assembly on varied geometries in cells suggests additional localization cues. We tested the hypothesis that lipid composition directs septin assembly through lipid packing properties. Lipid mixtures varying in lipid packing were designed by molecular dynamics simulations and incorporated onto supported lipid bilayers to measure septin adsorption in vitro. Septins strongly favor loosely-packed, disordered lipid bilayers but additional geometry cues act in conjunction with this membrane property. Introducing tighter lipid packing in cells disrupted septin structures in a curvature dependent manner, specifically limiting septin assembly and retention along flat regions of the plasma membrane. This work demonstrates that packing defects and geometry jointly regulate septin localization and highlights how multiple membrane properties are integrated to control organization of the septin cytoskeleton.

**Summary:** Localization of the septin cytoskeleton is controlled by regulatory factors, membrane curvature, and charge. In this study, changes to lipid composition that modulate lipid packing defects are found to impact septin assemblies in vitro and in cells.

## Introduction

The plasma membrane is an important surface for modulating many distinct protein assemblies from the cytoskeleton to the formation of condensates and the activation of signaling cascades (Snead et al. 2022; Banjade and Rosen 2014; Makushok et al. 2016; Martins et al. 2023; Aimon et al. 2014; Stone et al. 2017). Its lipid composition is tightly controlled, and the physical properties can vary over the course of the cell cycle (Vítová et al. 2021; Atilla-Gokcumen et al. 2014), due to environmental fluctuations (Gostinčar et al. 2019; Barrero-Sicilia et al. 2017; Cantarero et al. 2024), and at specific regions through changes to its curvature, charge, and lipid order (Fairn et al. 2011). Transmembrane and peripheral proteins act as both sensors and regulators of these conditions, contributing to the spatial organization of other cell factors and directing cell responses (Hofbauer et al. 2018; Lorent et al. 2017; Ballweg et al. 2020).

Septins are cytoskeletal proteins that can bind and assemble on the plasma membrane at specific locations and times to scaffold cell signaling events. In small eukaryotes, they are important players in cell division (Gladfelter et al. 2005), polarized growth (Spiliotis and McMurray 2020), and membrane organization (Barral et al. 2000; El Alaoui et al. 2025). In animals, these functions make septins critical for gamete formation (Al-Ali et al. 2024), tissue integrity (Kim and Cooper 2018), and neural architecture (Radler and Spiliotis 2022; Byeon et al. 2022). Septin assembly on membranes is regulated by other membrane-binding proteins (Longtine and Bi 2003; DeMay et al. 2009; Zheng et al. 2022), the presence of negatively charged lipids (Zhang et al. 1999; Bertin et al. 2010; Szuba et al. 2021), and through the emergence of shallow membrane curvature (Bridges et al. 2016; Lobato-Márquez et al. 2021; Beber et al. 2019). In turn, septins alter membrane properties by corralling certain species of lipids, limiting their diffusion (Pacheco et al. 2023), and are capable of deforming unsupported membranes (Tanaka-Takiguchi et al. 2009; Beber et al. 2019). Thus, septins both influence membrane properties and are themselves regulated by membrane association.

Septins have polybasic domains located along the N- and C-terminal extensions postulated to be important for septins’ preference for negatively charged lipids (Bertin et al. 2010; Cavini et al. 2024). However, N-terminal polybasic domains are located in the interface between septin subunits, making critical interface interactions and occluded from membrane binding (Mendonça et al. 2021; Cavini et al. 2021). Though evidence suggests GTP state may change septin conformation to expose these residues (Castro et al. 2020). Those on C-termini are often near a second membrane binding motif, an amphipathic helix (AH) domain. The AH domain is a well-conserved (Delic et al. 2024) stretch of amino acids required for septins’ preference for shallow membrane curvature (∼1 µm in diameter); in fungal and mammalian heterooctamers, this motif is located on the penultimate subunit on a long, flexible C-terminal extension (Cannon et al. 2019; Lobato-Márquez et al. 2021). Canonically, amphipathic helices detect highly curved membranes due to their ability to intercalate into the membrane using lipid packing defects, i.e. gaps between lipid headgroups exposing the hydrophobic fatty acid tails that are induced by membrane curvature (Cui et al. 2011; Antonny 2011; Bigay et al. 2005). However, changes to lipid packing via membrane curvature on a micron scale are predicted to be slight (Vanni et al. 2014; Badvaram and Camley 2023). Septins are nevertheless sensitive to subtle changes in curvature (Bridges et al. 2016; Cannon et al. 2019), raising the possibility that recognition of lipid packing defects may be one means leading to the emergence of curvature sensing in addition to other features of the membrane or through characteristics of septin assembly (Shi et al. 2023).

What is the possible contribution of lipid packing defects to septin membrane binding and assembly? Membrane curvature is one way lipid packing is altered but changing fatty acid tail saturation (Vanni et al. 2014) and the size of lipid headgroups (Vamparys et al. 2013) also change the characteristics of lipid packing defects. Saturated lipids exhibit tighter interactions between molecules and fewer openings between lipid headgroups while unsaturated lipids increase the number and size of lipid packing defects. In both cases, changes to the lipid composition can alter lipid packing independently of or in conjunction with membrane curvature, and these changes can alter AH domain localization in cells (Vanni et al. 2014; Lee et al. 2024). We hypothesized that lipid packing defects, as binding sites for the septin AH domain, may act as a critical cue for directing septin localization.

In this study, we used molecular dynamics simulations to design four lipid mixtures that vary in their lipid packing defect size or density and verified their differences using an in vitro membrane probe. By forming supported lipid bilayers using the different lipid compositions, we measured septin adsorption as a function of lipid packing and observed a strong preference for loosely packed, or more disordered, lipid mixtures. However, the data suggests that membrane geometry plays a role in addition to lipid packing that biases septin assembly on shallow membrane curvatures. Finally, we altered lipid packing in the model filamentous-growing fungus, *Ashbya gossypii*, and observed septin assembly depends on both lipid packing and membrane curvature in strong accordance with in vitro data. This study points to how both membrane geometry and lipid packing can be employed to recruit septins to a variety of different subcellular locations for their varied functions.

## Results

### Lipid composition modulates lipid packing of model membranes

Unlike most proteins with AH domains, septins’ curvature sorting behavior occurs on a micron scale rather than nanometer scale. On nanometer-scale curvature, defects increase substantially in size, density, and persistence (time they remain open) as membrane curvature increases (Vamparys et al. 2013; Gautier et al. 2018). In these cases, preferences arise primarily through the presentation of more binding sites rather than an increase in affinity (Hatzakis et al. 2009; Cui et al. 2011). However, septins exhibit a “goldilocks”-like preference, i.e. when presented with 0.5, 1, and 5 µm diameter SLBs, 1 µm SLBs have the highest septin adsorption, making septins’ reliance on lipid packing defects less clear cut (Bridges et al. 2016; Cannon et al. 2019).

To explore this discrepancy, we first designed a set of membrane compositions that could be used to test the role of lipid packing as a mechanism by which septins associate with membranes. Compositions span from the disordered to ordered phase while retaining a phosphatidylcholine motif on the headgroup and avoiding lipids that are likely to form a gel phase at room temperature, e.g. DPPC (Fig. 1A). To predict properties, membrane sheets of varying compositions (Fig. 1B) were built to a defined size via CHARMM Membrane Builder (Jo et al. 2009), and molecular dynamics simulations were run for 300 ns via GROMACS (Abraham et al. 2015), see example Fig. 1C. Simulations were analyzed in line with previous studies using the open-source software PackMem (Gautier et al. 2018) to retrieve the lipid packing defect constant, which allows for the comparison of the highest probability lipid packing defect size for each lipid composition. The largest defect constants were observed for Loose (L) and Moderate 1 (M1) compositions (14.9 Å^2^), which differed only by the presence of cholesterol. Moderate 2 (M2) and Tight (T) had markedly smaller lipid packing defect constants of 9.4 Å^2^ and 5.8 Å^2^, respectively (Fig. 1D). By using the number of defects counted in the PackMem analysis, we calculated the density of lipid packing defects for each membrane sheet and found it decreased with the degree of lipid packing (Fig. 1E). While L and M1 don’t differ in defect size, fewer defects arise in M1, presenting the opportunity to examine the impacts of both defect size and density on septin-membrane adsorption.

**Figure 1:**
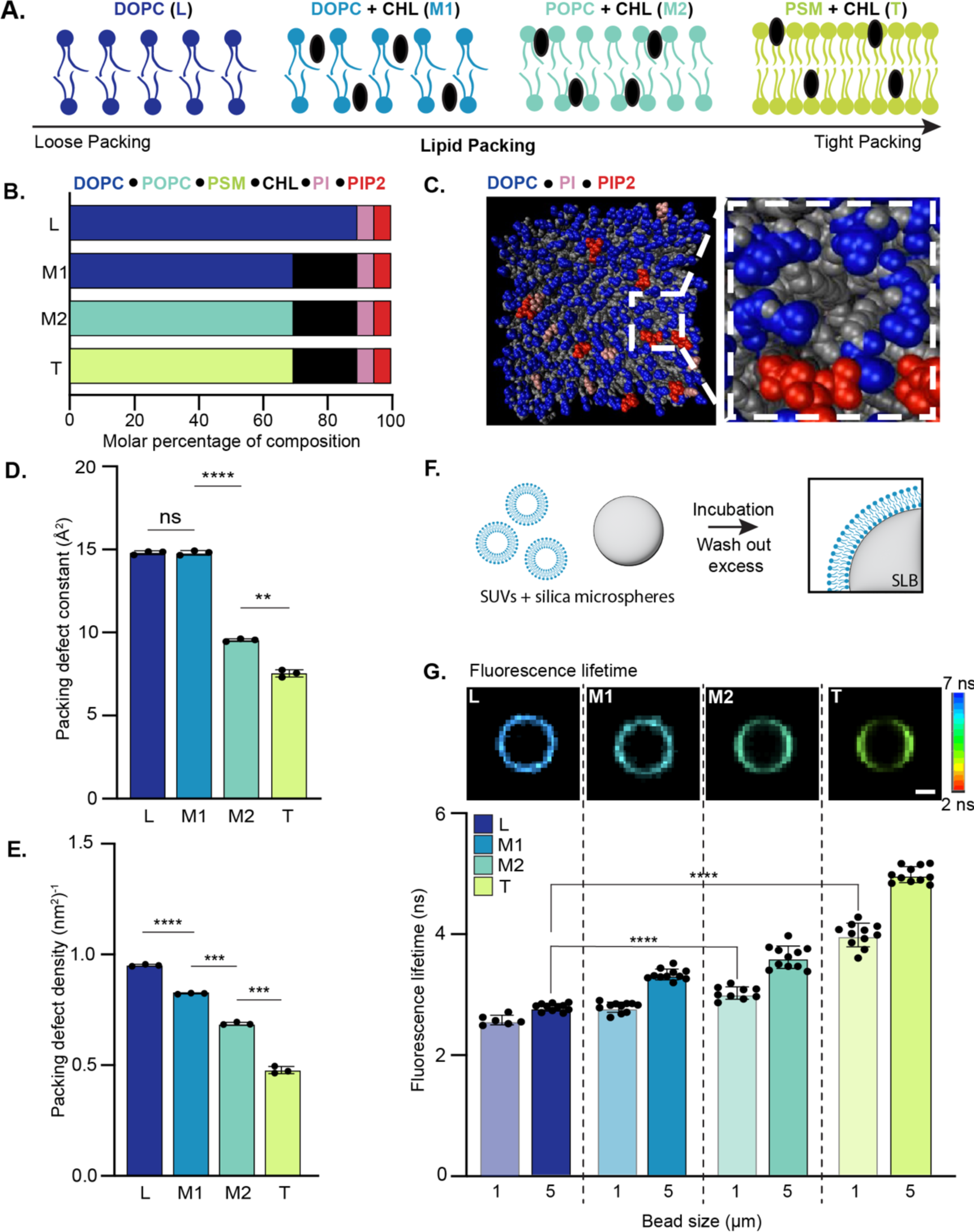
Lipid packing is modulated by lipid composition, both in silico and in vitro. (A) Cartoon representations of four lipid compositions with different degrees of lipid packing. **(B)** Full lipid compositions for L, M1, M2, and T. **Lipid abbreviations**: 1,2-dioleoyl-sn- glycero-3-phosphocholine (DOPC); Soy phosphatidylinositol (PI); 1,2-dioleoyl-sn- glycero-3-phospho-(1’-myo-inositol-4’,5’-bisphosphate) (PIP2); cholesterol (CHL); and 1- palmitoyl-2-oleoyl-glycero-3-phosphocholine (POPC). **(C)**. Snapshot of simulated lipid bilayer sheet and an example of a lipid packing defect where headgroups are in blue (DOPC), red (PIP2) and pink (PI). **(D)** Average lipid packing defect area for four lipid compositions, measured from simulations using open-source software PackMem. N = 3 independent simulations per composition. **(E)** Density of lipid packing defects measured for 11 x 11 nm membrane sheets of each lipid composition. **(F)** Schematic for SLB production: small unilamellar vesicles (SUVs) are incubated with silica microspheres of defined size. SUVs assemble onto the surface of microspheres and excess are washed away to produce bilayers. **(G)** In vitro reconstitution of lipid compositions onto spherical supported lipid bilayers (SLB) with FLIPPER-TR incorporated at 1 mol%. Fluorescence lifetimes measured by FLIM for each lipid composition on 1 µm and 5 µm SLBs. All differences between lipid compositions of the same curvature and between 1 and 5 µm SLBs of the same lipid composition are significant using unpaired t-test **** p<0.0001. Significance labels shown link to Fig. 4. Representative images of 5 µm SLBs, scale bar = 1 µm.

Characterizing lipid packing in cells and in vitro is challenging due to limited tools and the angstrom-level precision necessary to make direct measurements of lipid packing defect size, density, or depth empirically. To verify the predicted lipid packing behavior experimentally, we used FLIPPER-TR, a fluorescent molecule designed for reading membrane tension, as a proxy and incorporated it into model membranes on spherical supported lipid bilayers (SLBs) 1 and 5 µm in diameter (Fig. 1F). FLIPPER-TR reports membrane tension because it changes its conformation in response to pressure from the surrounding membrane, which is greater in tightly packed membranes than loosely packed ones (Colom et al. 2018). The conformational changes are detected as different fluorescence lifetimes, with tighter packing eliciting longer lifetimes than loosely packed membranes. The fluorescence lifetime of FLIPPER-TR was measured for each composition and as predicted, those lipid compositions with tighter packing measured in simulations have longer fluorescence lifetimes (Fig. 1G). Interestingly, fluorescence lifetimes measured for M1 are significantly longer than those measured for L, indicating FLIPPER-TR is capable of sensing differences in membrane properties not captured by the defect size constant measured in simulations. Notably, shorter fluorescence lifetimes (looser packing) were measured for FLIPPER-TR incorporated onto 1 µm SLBs compared to 5 µm despite very slight differences predicted for micron-scale curvatures (∼0.1 Å^2^, extrapolated from Vanni et al. 2014). In both cases, it’s likely FLIPPER-TR reports collective changes in defect density, depth, and persistence that alter pressure on the probe. These four lipid compositions provide a wide dynamic range of lipid packing defect sizes and densities with which to test the effects of lipid packing on septin assembly.

### Septin adsorption is sensitive to lipid packing

Septins possess an AH domain on the penultimate subunit, with two AH domains per heterooctamer (Fig. 2A). Previous work demonstrated that the septin AH domain is required for curvature-driven sorting to a preferred curvature of 1 µm in diameter (Cannon et al. 2019). Additionally, the AH domain is capable of detecting curvature as a standalone peptide in the absence of any other septin membrane-binding domains, suggesting septins use the AH to interact with membranes potentially through lipid packing defects (Cannon et al. 2019). To test the sensitivity of complete septin complexes to changes in lipid packing, we coated 5 µm diameter silica microspheres in each lipid composition identified in Fig. 1 and measured septin adsorption for each at steady state (Fig. 2B). 5 µm SLBs were used to assess septins’ sensitivity to lipid packing in the absence of the “ideal” membrane curvature (1 µm) while retaining substantial membrane association at higher salt concentrations (100 mM KCl).

**Figure 2:**
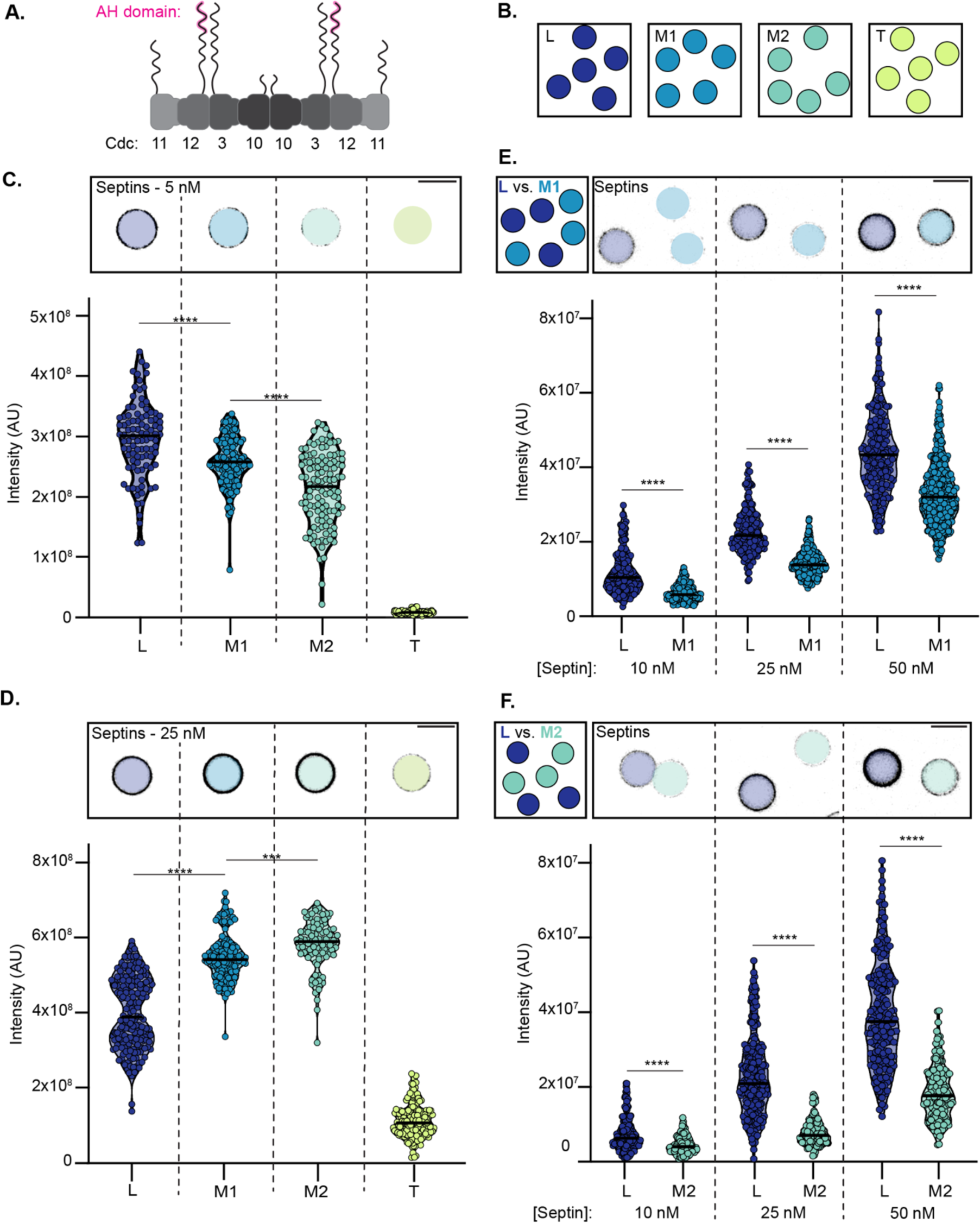
Septin assembly is sensitive to changes in lipid packing. **(A)** Schematic of *Saccharomyces cerevisiae* septin heterooctamer including location of amphipathic helix (AH) domain on Cdc12 C-terminal extension. **(B)** Schematic of experimental set-up where septin adsorption is measured on each lipid composition in separate reactions. **(C and D)** Septin adsorption on each lipid composition with representative micrograph in top panel at 5 nM **(C)** and 25 nM **(D)** septin. **(E and F)** Septins are incubated with two lipid compositions on the same SLB diameter with representative micrographs above each septin condition. **(E)** shows L and M1 in the same reaction and **(F)** shows L and M2 in the same reaction with 10, 25 and 50 nM septins for both. For C-F, black bars represent the mean, and significance was determined using an unpaired t-test, **** p < 0.0001.

We predicted that if septin AH engagement employs packing defects, septin adsorption would be higher on more loosely packed lipid compositions. Indeed, we see septin adsorption is strongly correlated with lipid composition at low septin concentrations (Fig. 2C). As lipid packing becomes tighter, septin absorption declines and is almost entirely impaired on the most tightly packed lipid composition T. We predicted this result based on the property of lipid packing, but septin localization to flat and concave membranes in *Ustilago Maydis* and *Aspergillus nidulans* is dependent on sphingolipid biosynthesis (Cánovas and Pérez-Martín 2009; Mela and Momany 2022). However, whether they bind directly to sphingolipids has been uncertain. Filipin staining of septin-tagged *U. maydis* shows primarily differential enrichment with some partial colocalization, and filipin treatment in *A. niger* is shown to be minimally disruptive to septin assemblies. It’s clear septins are heavily influenced by sphingomyelin-rich domains but our data suggest septins are unlikely to colocalize directly with them. Recent work by Goodchild et al. demonstrates using phase separated yeast polar lipid extracts that septins are occluded from ordered domains but align tightly to the boundary between disordered and ordered domains (in review). Combined, the data suggest the interphase boundary may offer favorable properties for septin binding, thus disruption of these phase-separated domains may be the cause of septin delocalization when sphingolipid synthesis is impaired in fungi.

For more moderate lipid packing compositions, we were somewhat surprised by the difference between L and M1 at low septin concentrations, as the two lipid compositions don’t vary in their lipid packing defect constant. Instead, they differ in defect density and perhaps other features as suggested by FLIPPER-TR. At near-saturating concentrations of septins (25 nM, Shi/Cannon 2022), septin adsorption is still largely absent on the tightest packing T SLBs, but septins’ preference shifts from L to M1 and M2 (Fig. 2D). This shift was surprising, but reminiscent of results seen in curvature sensing assays where high concentrations of septins lead to higher adsorption on non-preferred curvatures (e.g. 5 µm) compared to preferred (1 µm) when only one curvature is available for binding. This characteristic potentially indicates a greater ability for septins to pack more efficiently onto membranes of these compositions at high septin concentrations where septins are not limited in the bulk (Shi et al. 2023). However, it leaves septins’ absolute preference for lipid packing ambiguous.

In a cellular environment, septins are exposed to multiple cues simultaneously for localization and assembly, including membrane shape, regulatory factors, and a non-uniform plasma membrane. In such a competitive environment, is the membrane property of lipid packing capable of driving septin assembly to specific locations? To answer this question and approximate cellular membrane heterogeneity, we employed a “competition”-style experiment where two lipid compositions are present in equal proportions in the same reaction, “competing” for a common pool of septins.

We paired L with either M1 or M2 and measured septin adsorption at steady state for three septin concentrations, 10, 25 and 50 nM. In both competition cases, septins prefer L over the more moderately packed lipid composition at all three septin concentrations (**** p < 0.0001, Fig. 2E-F). These results suggest that lipid packing defects can bias septin localization, even at high septin concentrations, and demonstrates how situations where septins are limiting increases the effect of sorting preferences. However, this is a highly simplified environment compared to the complex context of a cell with many additional and potentially conflicting cues. Are septins able to sense, and be localized by, lipid packing in cells?

### Changes to fatty acid composition of cells alters septins in a geometry-dependent manner

Here we use *Ashbya gossypii*, a model filamentous fungus, to test the effects of changes to lipid composition on septins assembling on both flat and curved membranes. Multiple distinct and independent septin structures form in *A. gossypii*, including at the bases of branches (aligning with convex membrane curvature) and along the length of hyphae (flat membranes; Fig. 3A). On flat membranes, septins assemble into “inter-region (IR) rings” that are made of organized, bundled filaments (DeMay et al. 2009). Because many different septin assemblies are built in one cell within a continuous cytoplasm, these septin assemblies arise from a common pool of septins that “choose” between different membrane curvatures for assembly.

**Figure 3:**
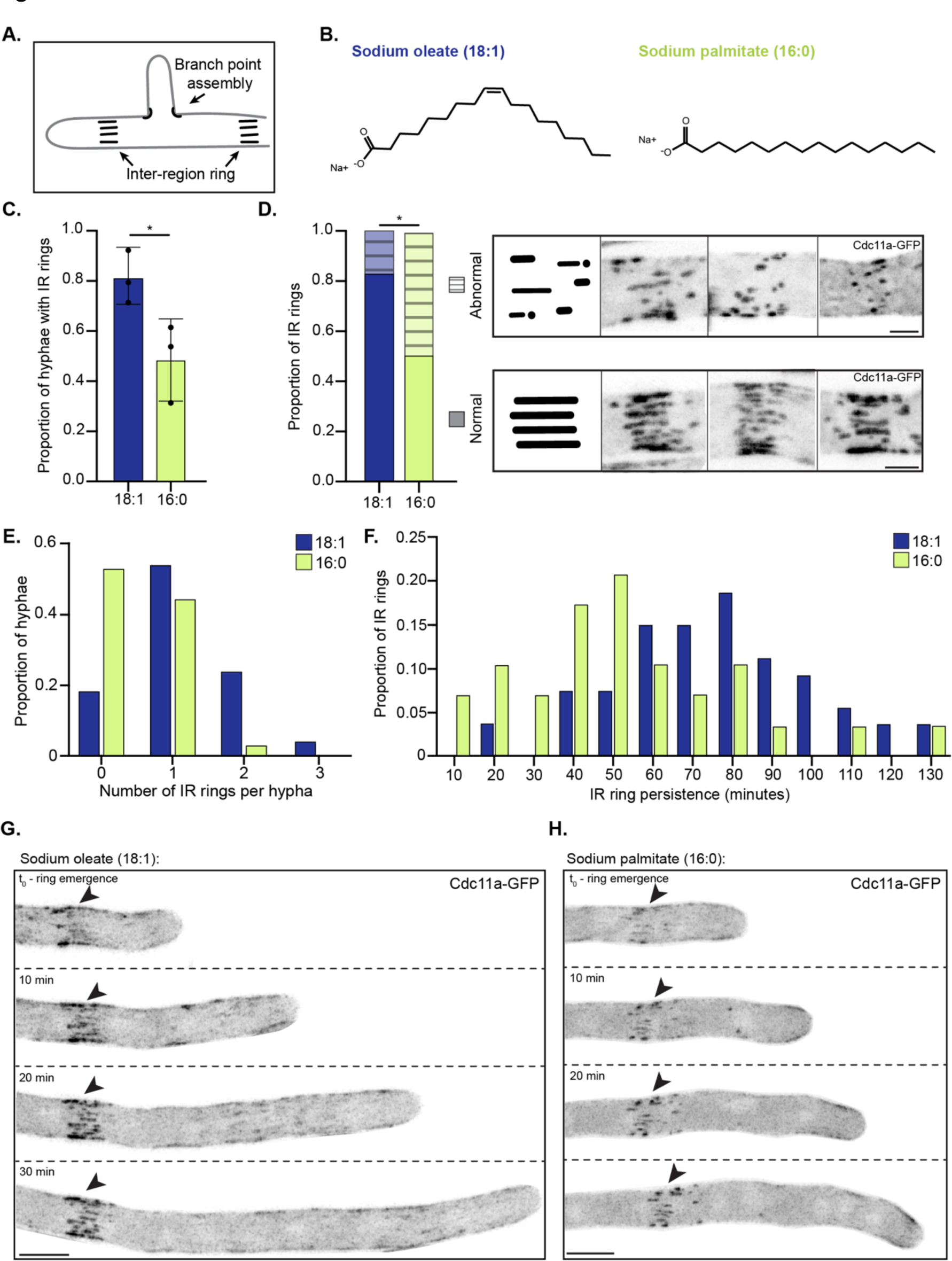
Changes to fatty acid composition in *Ashbya gossypii* alters septin assemblies along hyphae. **(A)** *A. gossypii* has septin structures at flat (along hyphae, IR rings) and curved regions of the plasma membrane (branch structures). **(B)** Fatty acids used in these experiments: sodium oleate (18:1) and sodium palmitate as (16:0). **(C)** Proportion of hyphae that build a septin IR ring post treatment in 140 minutes. N=76 hyphae for unsaturated and N=70 for saturated from three independent experiments per treatment. **(D)** Of those IR rings that are assembled under each treatment, the measured proportions that are “normal” and “abnormal” with cartoons and representative images of those classifications on the right. N=62 IR in unsaturated and N=33 in saturated fed cells. Scale bar for representative IR rings 2 µm. Significance by unpaired t-test * p<0.05. **(E)** Proportion of hyphae for each treatment that form 0, 1, 2, or 3 IR rings in the experimental window (140 minutes). **(F)** Histogram of the proportion of IR rings that persist for a given amount of time in the experimental window. **(G-H)** Representative hyphae assembling a septin IR ring over time for unsaturated **(G)** and saturated **(H)** fatty acid fed cells, corresponding to Movies 3 and 4. Scale bars 5 µm.

Does lipid packing impact the formation of these different assemblies? To address this, we wanted to perturb the normal balance of lipid composition in the plasma membrane, measure if and how septin assembly was impacted. To manipulate the fatty acid composition, cells were grown overnight, then shocked for 10 minutes with the fatty acid synthase inhibitor cerulenin to disrupt native fatty acid synthesis. The drug was replaced with minimal media containing fatty acid, either sodium oleate (unsaturated, 18:1) or sodium palmitate (saturated, 16:0; Fig. 3B) as well as a lower dose of cerulenin than the shock treatment to further prevent native fatty acid synthesis and encourage incorporation of supplemented fatty acids. We hypothesized that when palmitate was incorporated into the plasma membrane, it would tighten lipid packing, and if septin assembly is sensitive to packing as we saw in vitro, this would limit the formation of new septin assemblies. In contrast, incorporation of oleate should decrease lipid packing, opening or maintaining lipid packing defects to promote assembly.

After feeding with different fatty acids, cells were immediately plated onto an agarose pad and imaged every 10 minutes to observe the assembly process of septin structures. In each movie, we calculated the proportion of hyphae that formed IR rings over 140 minutes and found that on average, 77% of hyphae (N=76) formed an IR ring when fed unsaturated sodium oleate and only 48% of hyphae (N=70) formed when fed saturated sodium palmitate (Fig. 3C). In addition, the IR rings were not formed equally well in terms of size and organization. For the first IR ring formed in each hypha, we measured the proportion with “normal morphology” (well-organized bundled filaments) and those with abnormal morphology (fragmented or punctate; Fig. 3D). For unsaturated fatty acid fed cells, an average of 82% of IR rings (N=62) had normal morphology while only 51% (N=33) looked normal when fed saturated fatty acids. Next, we measured the number of IR rings formed per hypha and found cells fed unsaturated fatty acids were more likely to produce more than one IR ring in the same amount of time (Fig. 3E, N=76 hyphae).

However, we measured the length of new growth for each hypha and observed more growth in unsaturated fed cells compared to saturated, which could account for the increase in number of IR rings, as they continue to be laid down at a defined interval (every 40-65 µm) in healthy cells (Fig. S1B).

Finally, we measured the persistence of IR rings under both treatments (Fig. 3F). IR rings are the precursor for septa, a barrier of plasma membrane and cell wall used to close off parts of the hyphae in response to age and environmental stress. We measured the time in existence of an IR ring from formation to the end, including both those transitioning into a septum and those that appear to “dissolve” or lose septin intensity from the membrane without making a septum (example in Fig. S1A, see movies 1 and 2). There is a clear shift in the persistence of the first IR ring formed depending on treatment, with the mode lifetime of IR rings under saturated treatments being 50 minutes (N=33 IR) and the mode for unsaturated being 80 minutes (N=62 IR). It should be noted that these measurements are significantly under-counted as we cannot account for those IR rings still present at the end of each movie, which is a little under half in saturated fed hyphae (41%) and the majority in unsaturated fed hyphae (81%). In Fig. 3G-H, we show representative stills of the formation of IR rings over time for each treatment (see corresponding movies 3 and 4), which is characterized by an initial structure of thin filaments that becomes more bundled over time, however, in saturated fatty acid fed cells, these structures often assemble with only short septin filaments and puncta. Interestingly, short assemblies that do form can grow brighter over time, suggesting lateral septin interactions remain unimpeded. These deficiencies in assembly likely arise from reduced defect numbers (fewer AH binding sites) and weaker septin anchoring due to changes in defect depth or persistence.

We also wanted to analyze septin assembly at sites of curvature but surprisingly few branches formed during both treatments. Those that did form predominantly emerged later in the movies when the actual lipid composition may be more heterogeneous and usually in regions preceding or right at the start of new growth (Movie 5). In both conditions, with the exception of one hypha in a saturated sample, septins were enriched at convex curvature at branch points as in untreated cells (Fig. S1C). Taken together and combined with in vitro data, these results indicate that septins are intrinsically sensitive to the membrane property of lipid packing and that it can be used to modulate their assembly in cells. However, these changes appear not to be sufficient to affect septin localization at shallow curvature at branch points, indicating there might be an additional feature at these sites that septins are recognizing to cue their assembly.

### Composition-induced lipid packing and geometric cues combine to direct septins to the preferred curvature

We have demonstrated that septins are sensitive to lipid packing similarly to other curvature sensing proteins, but while changes to lipid packing in cells is sufficient to disrupt assembly on flat membranes, binding to curved membranes is unaffected. Septins’ preferred membrane curvature of 1 µm is predicted to vary only slightly from 5 µm diameter bilayers, yet FLIPPER-TR detects significant differences between 1 and 5 µm beads for all lipid compositions. To test if lipid packing is the primary feature recognized by septins on curved membranes, we modified the competition assay to include both different curvatures and lipid compositions in the same reaction.

In each well, 5 µm spheres are coated in L lipid composition so that the non-preferred membrane curvature (5 µm) has the preferred lipid composition (L). These L-5 µm SLBs are then paired with preferred curvature (1 µm) and less optimal tight lipid packing (either M2 or T lipids). In both cases, we expect L-5 µm to have the loosest packing (by size and density) compared to 1 µm beads of either composition. This is supported by FLIPPER-TR results from Fig. 1F that shows L-5 µm SLBs had the shortest lifetime, significantly shorter than M2-1 µm and T-1 µm (**** p<0.0001). We hypothesized that if lipid packing is the driving force of septin curvature sensing, septins would accumulate on L-5 µm, even if 1 µm beads of either M2 or T were present. Alternatively, if septins use a mechanism in addition to lipid packing to sense a specific membrane curvature, we expected septins to show the highest adsorption on the preferred curvature, despite more tightly packed lipids.

Strikingly, when L-5 µm and M2-1 µm are incubated together, septins show the highest adsorption on M2-1 µm SLBs (Fig. 4A). Septins still coat 5 µm spheres evenly, but the mean adsorption on M2-1 µm is ∼1.5 times greater than L-5 µm. In this case, the ∼5 Å^2^ difference in defect constants as well as the lower expected density of lipid packing defects between L and M2 via simulations wasn’t enough to override septins’ curvature sorting behavior. But what if the lipid composition is very tightly packed? We tested this by incubating septins on L-5 µm and T-1 µm, the tightest packing lipid composition composed primarily of sphingomyelin and cholesterol. In this case, septins were predominantly unable to bind the tightly packed 1 µm SLBs and instead showed high adsorption onto 5 µm beads of L lipid composition (Fig. 4B).

**Figure 4:**
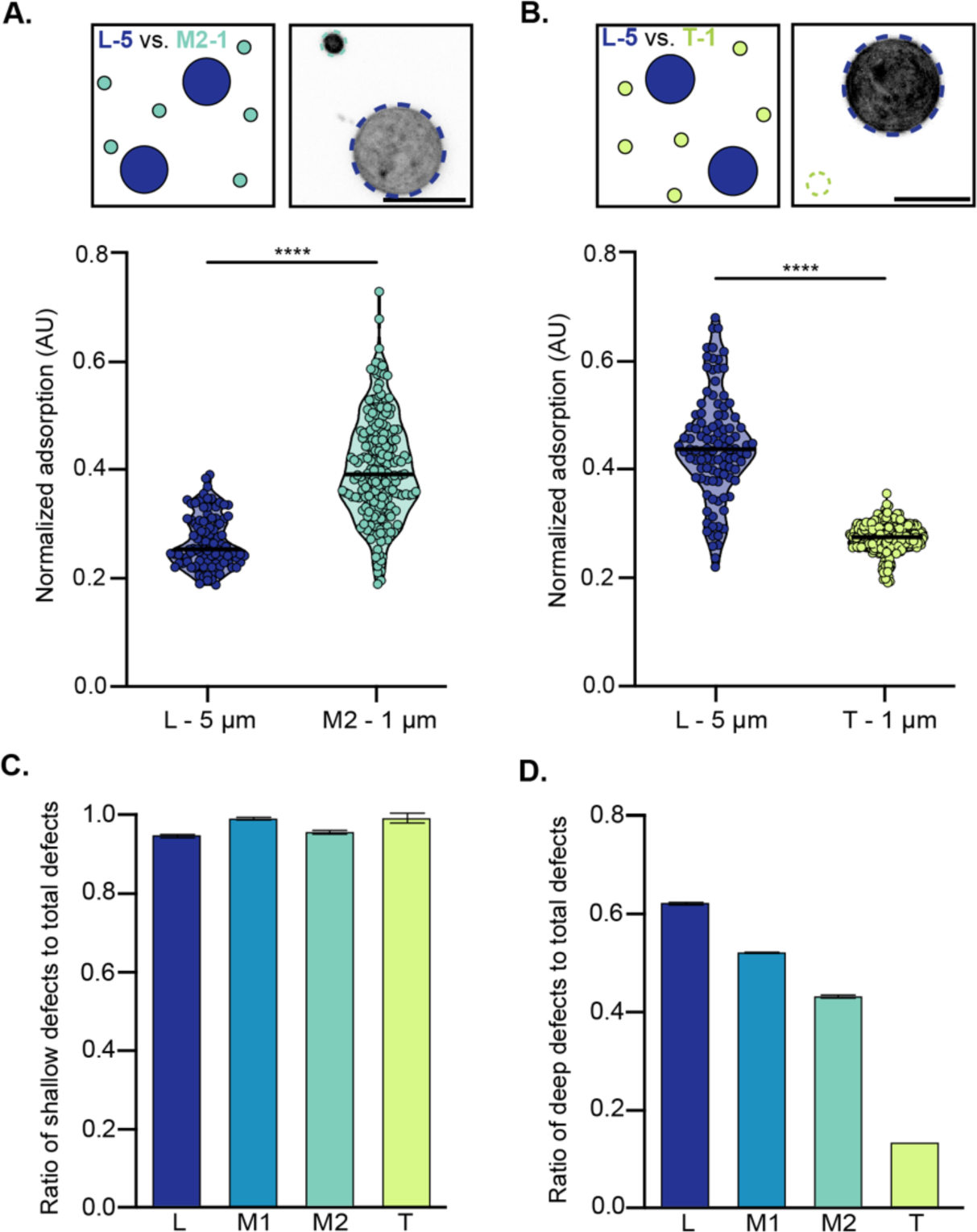
Looser lipid packing is not sufficient to drive septins to more shallow membrane curvatures. **(A-B)** Combination of shape and membrane composition in competition with 5 nM septins. Septin adsorption normalized for membrane surface area is plotted with representative micrographs in top panel for L-5 µm incubated with M2-1 µm **(A)** and L-5 µm incubated with T-1 µm **(B).** Scale bars 5 µm. Significance by unpaired t-test **** p<0.0001. **(C-D)** Ratio of the number of shallow defects **(C)** and deep defects **(D)** to the total number of defects measured from molecular dynamics simulations of membrane sheets, measured through PackMem. For T, two of the three simulations failed to report any deep defects, thus only the proportion for one simulation is reported in **D.**

Is the rate of adsorption onto the loosely packed lipid composition so high that it out- competes the alternative mechanism of curvature sensing or is T too tightly packed and blocking septins from binding? We returned to molecular dynamics simulations of membrane sheets and took a closer look at the types of defects counted in each simulation. Defects are categorized as “shallow” defects that only expose fatty acid tails close to the surface of the membrane and “deep” defects that extend ≥ 1 Å below the central glycerol carbon. Each contiguous packing defect counted (total defects) may be made up of multiple shallow and/or deep defects, thus the sum of deep and shallow defects doesn’t add up to the total number of defects counted, but we can use the relative ratios of different depths to make inferences about the landscape an AH domain may encounter. Interestingly, the ratio of shallow defects to total defects doesn’t change substantially as lipid packing changes and is close to 1, i.e. most packing defects contain a shallow region (Fig. 4C), but the ratio of deep defects to total defects declines according to changes in lipid packing (Fig. 4D). Thus, it’s possible septins require lipid packing defects of a minimum size and/or depth to assemble onto the “preferred” membrane curvature (Cui et al. 2011; Fu et al. 2024).

This in vitro result complements what we observed in *A. gossypii* where lipid feeding experiments that block septin assembly on flat membranes do not affect septin assembly on their preferred membrane curvature. It seems likely in Fig. 3 experiments that *A. gossypii* doesn’t allow its plasma membrane to become very tightly packed, or simply the rate of sodium palmitate adsorption and subsequent incorporation into lipid synthesis is too slow to reach the degree of lipid packing necessary to block septin assembly at the optimum curvature seen at branch sites. Taken together, our data indicate that the property of lipid packing is sufficient to cue septin assembly independently of membrane geometry but suggests an additional mechanism that biases septin assembly on curved membranes, contributing to septins’ ability to sense micron-scale membrane curvature (Fig. 5).

**Figure 5:**
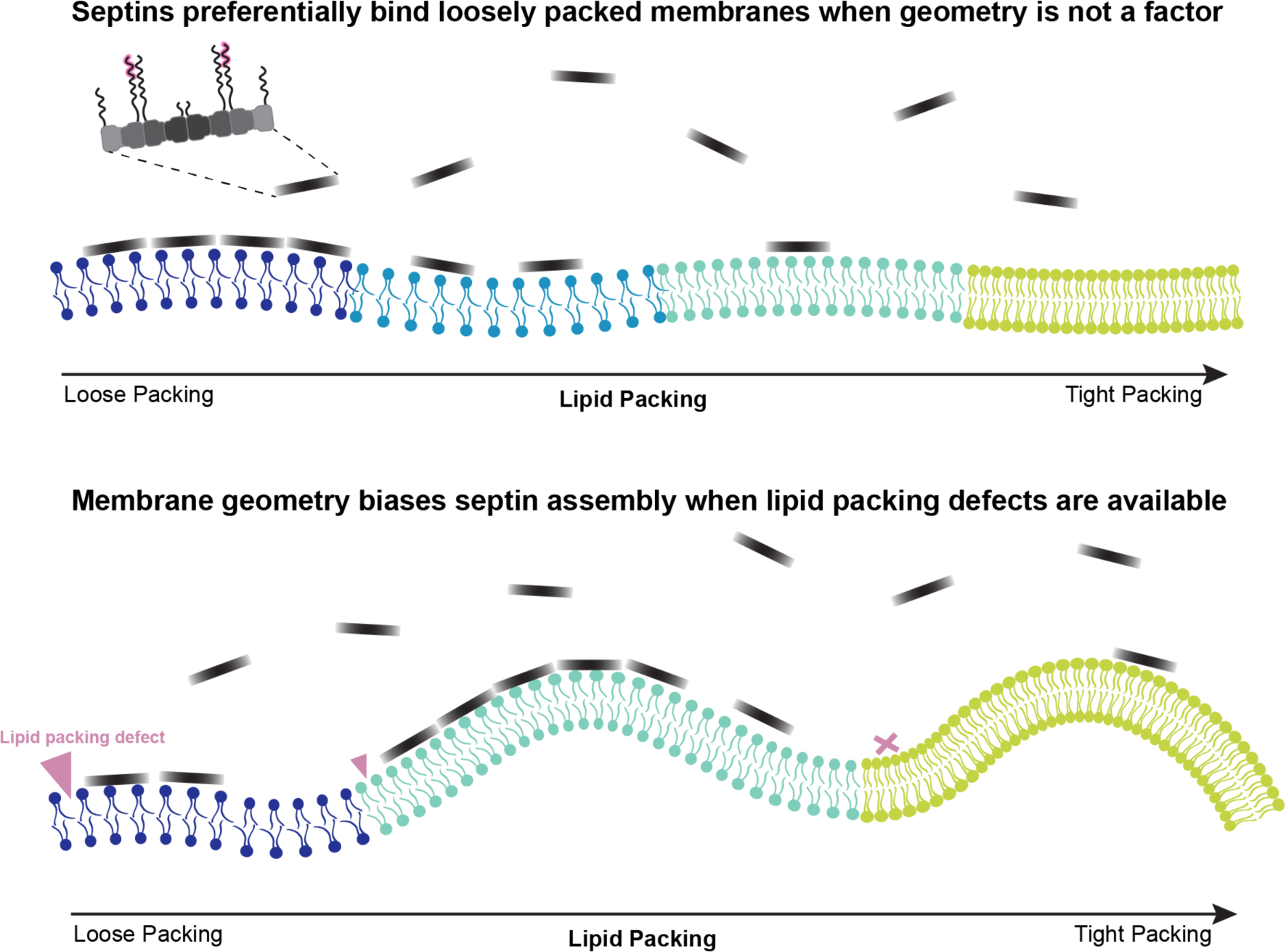
Septin adsorption is directed by lipid packing defects that provide binding pockets for the amphipathic helix domain on Cdc12. When membrane geometry is uniform, septins preferentially bind loosely packed lipids, but when more tightly packed lipid compositions are paired with septins’ preferred membrane geometry, septins are biased towards the ideal with geometry with lipid packing defects of a minimum size and depth.

Future research is necessary to identify the nature of this curvature-dependent mechanism, which may arise through multiple means, such as 1) rates of septin association, 2) septin oligomerization, 3) flexibility of the oligomer or filament, and 4) septin-induced membrane properties. Previous research has highlighted that septins are near the physical limit for measuring local curvature on a micron scale based on their curvature-dependent association rates (Badvaram and Camley 2023). In their model, they see greater sensitivity to local curvature than to lipid density, also pointing to alternative modes of curvature sensing. Notably, other curvature sensors that lack AH domains or combine AH and BAR domains also exhibit defined curvature preferences (Simunovic et al. 2015). The origins of these preferences are still unclear, but the ability to self-assemble is a common thread among BAR domain containing proteins, septins, and another micron-scale curvature sensor, SpoVM (Gill et al. 2015). Both septins and SpoVM require oligomerization to sense their preferred curvature (Kim et al. 2017; Shi et al. 2023), and septin assembly becomes more cooperative on preferred membrane curvature (Cannon et al. 2019). This could result from conformational shifts that promote end-on interactions, due to the flexibility of the septin heterooligomer (Mendonça et al. 2021), or flexibility of septin filaments which can be regulated by septin composition (Khan et al. 2018). Lastly, because a minimum defect size or density is required and septins are known to induce local membrane curvature (Beber et al. 2019), it is possible septins may alter membranes after binding. For example, the physicochemical properties of a membrane may change in a geometry-dependent manner, such as through long-range interactions or altering lipid packing defect distribution that could facilitate septin recruitment (Simunovic et al. 2017; Reynwar et al. 2007). In conclusion, lipid packing defects are sufficient to localize septins in a geometry-independent manner but act in conjunction with geometry-dependent cues as part of a suite of mechanisms that promote micron scale curvature sensing.

## Movie legends

**Movie 1**: *Ashbya gossypii* with fluorescently labeled Cdc11a-GFP imaged every 10 minutes via laser scanning confocal treated with sodium palmitate (16:0). This movie shows how IR rings form and dissolve under this treatment. Stills are shown in Figure S1A. Yellow arrows follow the IR ring progression.

**Movie 2**: *Ashbya gossypii* with fluorescently labeled Cdc11a-GFP imaged every 10 minutes via laser scanning confocal treated with sodium palmitate (16:0). This movie shows another example of how IR rings form and dissolve under this treatment, though this movie highlights how this happens in a long, fast-growing hypha. Yellow arrows follow the IR ring progression.

**Movie 3**: *Ashbya gossypii* with fluorescently labeled Cdc11a-GFP imaged every 10 minutes via laser scanning confocal treated with sodium oleate (18:1). This movie corresponds to Figure 3G, showing a representative hypha forming a “normal” IR ring. Dark blue arrows follow the IR ring progression.

**Movie 4**: *Ashbya gossypii* with fluorescently labeled Cdc11a-GFP imaged every 10 minutes via laser scanning confocal treated with sodium palmitate (16:0). This movie is representative of how “abnormal” IR rings form under this treatment, corresponding to Figure 3H. In this case, the IR ring persists to the end of the movie. Yellow arrows follow the IR ring progression.

**Movie 5**: *Ashbya gossypii* with fluorescently labeled Cdc11a-GFP imaged every 10 minutes via laser scanning confocal treated with sodium oleate (18:1). This movie shows a hypha that forms two IR rings and two branches under this treatment. Thick dark blue arrows follow the IR ring progression while thinner arrows follow septin enrichment at the branch sites.

## Materials and Methods

### Molecular dynamics simulations

Molecular dynamics simulations of flat lipid bilayers were initiated using Charmm Gui Membrane Builder (https://www.charmm-gui.org/). Bilayers were built to have 160 lipids per leaflet in 100 mM KCl, and at 298.15 K (25°C), to approximate experimental conditions. Simulations were equilibrated for a total of 1.5 µs before production of 300 ns. Three independent replicates were performed through random seed generation. Analysis of lipid packing defects was run using the open-source software Packmem (https://packmem.ipmc.cnrs.fr/) with sampling every 100 ps.

### Septin protein purification and quality check

To purify yeast septin proteins, BL21 (DE3) cells were transformed using a duet expression vector (Bridges 2016) under ampicillin and chloramphenicol selection. Transformants were cultured in terrific broth (Gibco) to an OD600 between 0.6-0.8 and induced with 1 mM isopropyl *β*-d-1-thiogalactopyranoside. Induced cultures were grown at 18°C for 18 hours before collecting cell pellets by centrifugation at 7k RPM for 15 minutes. 1L cell pellets were frozen at -80°C and thawed in 35 mL lysis buffer (1 M KCl, 50 mM HEPES pH 7.4, 1 mM MgCl2, 10% glycerol, 1% Tween-20, and 1 EDTA-free protease inhibitor tablet (Pierce), 20 mM Imidazole, 1 mg/mL lysozyme, and 5 units RQ1 DNase in ice water with intermittent vortexing. Cell mixture was then homogenized in a 40 mL glass homogenizer until smooth and sonicated for a total of 10 minutes with microtip probe sonicator at 70% amplitude (Branson). Lysate was clarified by centrifugation in a SS-34 rotor at 20k RPM for 30 minutes. Septin purification was performed on an AKTA GO FPLC by loading clarified lysate onto a Cytiva 5 mL HisTrap HP column equilibrated with lysis buffer, washed with 50 mL of Wash buffer (1 M KCl, 50 mM HEPES pH 7.4, 40 mM imidazole) and eluted with 50 mL of Elution buffer (300 mM KCl, 50 mM HEPES pH 7.4, 300 mM Imidazole). Eluent was concentrated to 500 uL before running over a Superdex 200 Increase 10/300 size exclusion column in SEC buffer (700 mM KCl, 50 mM HEPES pH 7.4, 1 mM *β*-mercaptoethanol). Higher salt helps improve purity. Septins were dialyzed into septin storage buffer (300 mM KCl, 50 mM HEPES pH 7.4, 10% glycerol, 1 mM *β*-mercaptoethanol) for 24 hours in two steps (Slide-A-Lyzer G2; 20,000 molecular weight cut-off (Thermo Fisher Scientific)). In step two of dialysis, 60 µg of Tobacco etch virus protease (Sigma) was added to cleave the 6×- histidine tag on Cdc12. After dialysis, protein was run over a second column of cobalt resin to remove the protease, poly-histidine tag, and other contaminants. Septins were labeled with SNAP Surface 488 (New England Biolabs) overnight with 2x the protein concentration of dye. Protein purity was determined via SDS-PAGE and protein concentration by Bradford assay. Protein quality was assessed by performing a curvature sensing assay with 1 and 5 µm supported lipid bilayers and by observing filament formation at 100 mM KCl on PEG-passivated coverslips.

### SUV preparation

Small unilamellar vesicles (SUVs) of different lipid compositions (Avanti Polar Lipids) were prepared to make SLBs. For all compositions, lipids were mixed in chloroform by mol% listed in Fig. 1B with the addition of 0.01% Eggliss Rhodamine PE or 18:0 Cy5 PE, dried by a stream of nitrogen gas to form a lipid film, then dried under vacuum overnight. For experiments testing the relative packing of lipid compositions in vitro, a final concentration of 2 mol% FLIPPER-TR (Spirochrome) dissolved in fresh DMSO was added to lipid mixtures before drying. Lipids were resuspended in supported lipid bilayer buffer (300 mM KCl, 20 mM HEPES pH 7.4, and 1 mM MgCl2) for a final concentration of 5 mM over 30 minutes at 37°C with vortexing every 10 minutes. In 2 mL amber vials, hydrated lipids were sonicated for 10-15 minutes until clarified.

### Spherical supported lipid bilayers (SLB)

SUVs were adsorbed onto silica microspheres (Curtis et al. 2022) of various curvatures by mixing 50 nM lipids with microspheres for 1 h on a roller drum at room temperature. Unbound lipid was washed away by pelleting lipid-coated beads at the minimum force required to pellet each bead size (for diameters 1 µm = 2300 RCF and 5 µm = 300 RCF) and washed using pre-reaction buffer (33.3 mM KCl and 50 mM HEPES, pH 7.4) four times, then diluted to a total surface area of 5 mm^2^ in reaction buffer (33.3 mM KCl, 50 mM HEPES pH 7.4, 150 µg/mL beta casein, 0.1% methyl cellulose). For reactions with either multiple lipid compositions or multiple SLB diameters, total surface area was kept constant, with equal surface area for each composition/diameter. For incubation with septins, 25 uL of septins in storage buffer were mixed with 75 uL reaction buffer and gently mixed in home-made wells on PEGylated glass coverslips. Equilibration occurs over 2-2.5 hours depending on the lipid composition.

### Fluorescence microscopy

FLIM: To measure the relative lipid packing of different compositions in vitro, we used the membrane tension probe FLIPPER-TR. Fluorescence lifetime imaging microscopy (FLIM) was performed on the Leica SP8X Stellaris (Neuroscience Microscopy Core) using a 63X oil objective, WLL excitation at 488 nm, 70% laser power, 20 MHz pulses, and emission filter at 550-650 nm. Single z slices were taken along the midplane of the SLBs. Analysis was performed with built-in Leica software, using a two-term exponential decay function with correction for IRF.

Confocal: To measure septin adsorption, each septin/SLB mixture was incubated in a plastic chamber glued to a polyethyleneglycol (PEG)–passivated coverslip until equilibrium (∼2.5 hours, Curtis and Vogt et al., 2022). Beads were imaged using a Zeiss 980 laser scanning confocal under Airyscan MPLX 4Y mode with a 63x oil objective. For analysis, airyscan processing at a defined threshold was applied to all images, then septin intensity values corresponding to lipid masks were collected by Arivis. When two SLB sizes were present, images were collected on a Nikon Ti-2 with Yokogawa CSU-W1 spinning disk using 100x silicone oil objective. For comparing between different diameter SLBs, controls were collected of each lipid composition on both bead sizes. One composition was chosen arbitrarily to be the “normal” values, and the sum intensities of the lipid channel were normalized to their corresponding size of that composition. Normalization to both compositions was tested to ensure results were not biased by this step. Septin adsorption was then calculated by dividing the septin sum intensity values over the normalized lipid sum intensity values to control for membrane surface area. All statistical analyses were performed on Graphpad Prism using unpaired parametric t-tests.

### Ashbya gossypii culture. fatty acid feeding experiments, and imaging

Fatty acid feeding experiments were adapted from previous work (Makarova et al. 2020). *Ashbya gossypii* spores were germinated in full medium 25°C for 22 hr and harvested by centrifugation in a swinging bucket rotor at 300 rpm. Full media was replaced with minimal media containing 100 µM cerulenin, and cells were shaken at 25°C 10 minutes. This media was then replaced with minimal media containing 20 µM cerulenin and 200 µM of either sodium palmitate or sodium oleate (Thermofisher). To prepare a concentrated stock solution of fatty acids, either sodium palmitate or sodium oleate powder was added to minimal media and dissolved through sonication for 5-10 minutes in a bath sonicator until fully emulsified before being diluted into treatment media. Cells were then harvested again by centrifugation and mounted on minimal media pads containing 2% agarose for immediate live imaging. Full Z stacks were acquired from Zeiss 980 laser scanning confocal under Airyscan MPLX 4Y mode using a 40X water objective, and 488 nm excitation at 0.8% laser power every 10 minutes for 140 minutes. All statistical analyses were performed on Graphpad Prism using unpaired parametric t-tests.

## Acknowledgements

We would like to thank Adai Colom for his discussions on the use of FLIPPER-TR for supported lipid bilayers. Fluorescence Lifetime Imaging Microscopy was performed at the UNC Neuroscience Microscopy Core (RRID:SCR_019060), supported in part by funding from the NIH-NINDS Neuroscience Center Support Grant P30 NS045892 and the NIH-NICHD Intellectual and Developmental Disabilities Research Center Support Grant P50 HD103573. This work was funded by NIGMS under award T32 GM119999, NIGMS award 1F31GM151820-01A1, NIH grant R01GM130934, Alfred P. Sloan Foundation grant G- 2021-14197, and HHMI.

---

**Figure S1:**
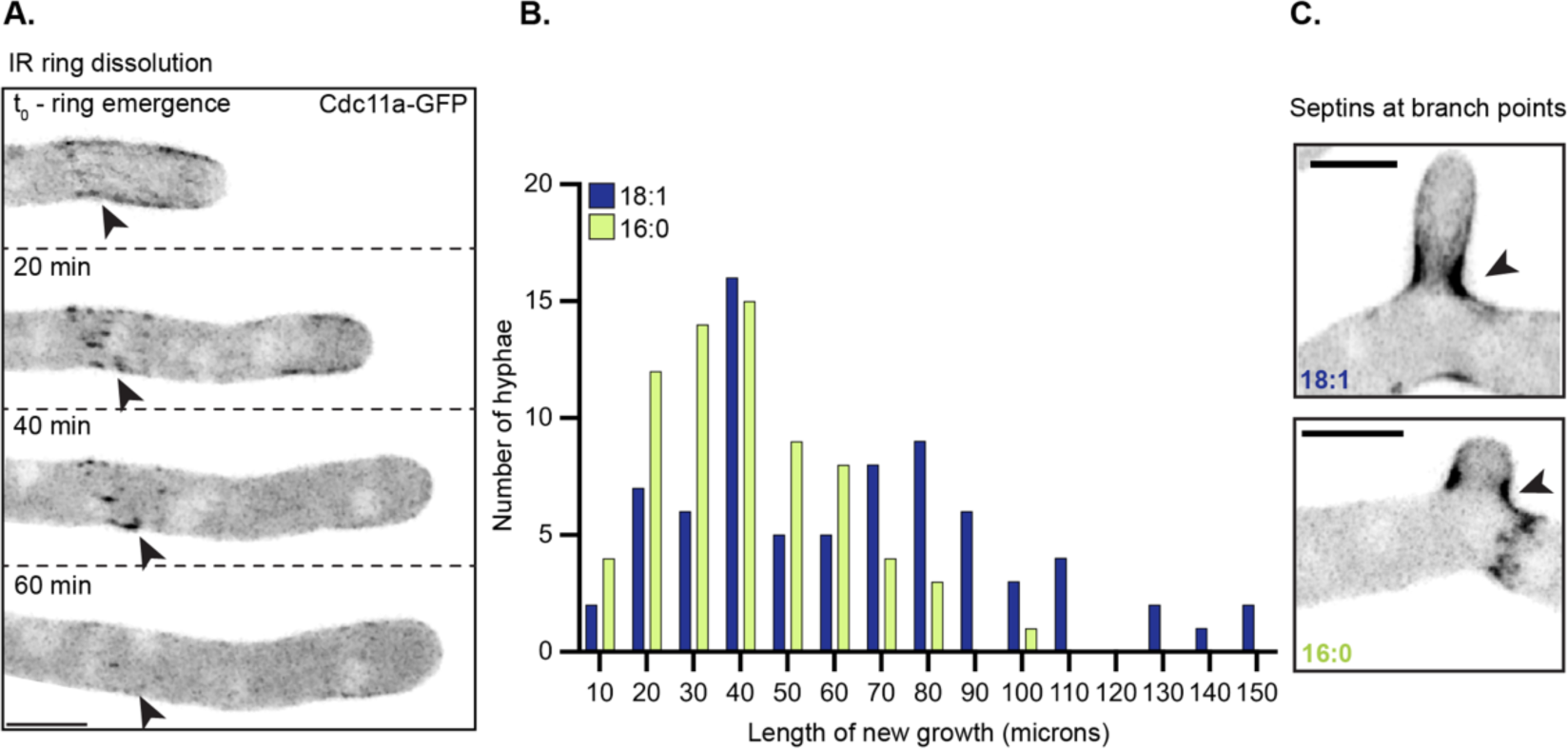
Fatty acid effects on new growth and septin assemblies. **(A)** An example of an IR ring emerging and disappearing over the course of 60 minutes in sodium palmitate (16:0) fed cells, also shown in Movie 1. Scale bars 5 µm. **(B)** Histogram showing the distribution of new growth binned in 10 micron increments and the number of hyphae for each treatment that fall into each category. **(C)** An example from each treatment of septins enriched at branch points in *A. gossypii*.

